# Technical insights into fluorescence lifetime microscopy of mechanosensitive Flipper probes

**DOI:** 10.1101/2022.09.28.509885

**Authors:** Chloé Roffay, Juan Manuel García-Arcos, Pierrik Chapuis, Javier López-Andarias, Falk Schneider, Adai Colom, Caterina Tomba, Ilaria Di Meglio, Valentin Dunsing, Stefan Matile, Aurélien Roux, Vincent Mercier

## Abstract

Measuring forces within living cells remains a technical challenge. We developed hydrophobic mechanosensing fluorescent probes called Flippers, whose fluorescence lifetime depends on lipid packing and can report on membrane tension. Here, we describe technical optimization of the probe imaging, and diverse characterizations in various biological and in vitro systems. We provide a guideline to measure biophysical parameters of cellular membranes by FLIM microscopy with Flipper probes, providing evidences that flippers can report long range forces in cells, tissues and organs^i^.

## Introduction

Measuring physical forces in biological material is of increasing interest, but remains technically challenging[1]. The possibility of measuring such forces through microscopy imaging presents the advantages of being less intrusive than many mechanical methods, making use of the universally available microscopy equipment in research labs. Many fluorescence tools have been developed in this regard, in particular Forster Resonance Energy Transfer (FRET) biosensors, in which the elongation of a linker between two FRET fluorophores under force reduces FRET efficiency. The FRET couple and the linker can be composed of fluorescent proteins connected by a peptide chain, all being easily expressed in cells using standard molecular biology techniques. This has been developed to measure forces at focal adhesions, and has been extended to measure force between membranes and actin[2], the cytoskeleton and the nucleoskeleton[3] or under shear stress[4]. However, these biosensors have several limitations, namely that the force range is very small, and thus require the use of several chimeric constructs with different linkers. Also, it requires the force sensor to be bound to proteins "handles" onto which the force is applied and finding the right combination of interacting domains that will not detach under force remains challenging.

Another strategy has been to design small chemicals whose fluorescence properties will change under force variation. For example, molecular rotors report change in membranes viscosity by variations on their fluorescence lifetime which is measured by Fluorescence Lifetime Imaging Microscopy (FLIM) [5,6]. One of the challenges has been to expand the list of forces that can be measured, as some of the forces are exerted within materials, and not between materials. We have designed, synthesized and characterized small-molecules called FLIPPERS that can report changes of membrane lateral forces due to lipid packing variations through changes of their conformation and hence of their fluorescence properties[7]. The high sensitivity of Flippers allows them to report membrane tension variations by changes of the lifetime of their excited state or fluorescence lifetime[8], upon osmotic shocks[9,10] or to investigate cellular processes such as lysosomal exocytosis[11]. The flipper probes were used to study TORC2[12,13], internalized to study nuclear membrane properties[14], in the context of 3D migration[15] or membrane properties at nucleation zone[16]. Moreover studies investigated membrane tension in mature adipocytes[17], revealed distinct lipid microdomains in young epidermal cells in the meristem[18], probed the role of ether lipids and sphingolipids in the early secretory pathway[19], determined the effects of glycolysis inhibition on membrane order[20], compared membrane tension within crypt-villus structures during intestinal development[21]. Over the past six years, we designed various derivatives targeting different organelles[22] (plasma membrane, endoplasmic reticulum, mitochondria and lysosome/late endosome). We also developed probes with different biochemical and/or photochemical properties[6] (HaloFlipper, HaloPhotoFlipper, HydroFlipper). We identified fluorescence lifetime imaging (FLIM) as the best way to report the conformational changes of Flipper probes with lipid packing because the amplitude of lifetime variation is higher than the change of fluorescence intensity or wavelength shift.

However, measuring membrane tension using FLIM of Flippers remains technically challenging. This method paper provides technical and practical information to make the use of Flipper probes for membrane tension measurements more accessible and reproducible.

## I-Basics of Flipper probes

Four commercialized Flipper probes with identical mechano-chemistry principle target different membranes in cells: Flipper-TR for plasma membrane, ER Flipper-TR for endoplasmic reticulum, Lyso Flipper-TR[22] for late endo/lysosomes and Mito Flipper-TR for mitochondria. The common chemical structure of Flipper probes (see Fig 1a) is composed of two dithienothiophenes (DTT) fluorescent groups that can rotate around the carbon bond that links them together. Methyl groups have been added at specific places to ensure that the two DTT groups are twisted out of co-planarity in the ground state but can still be planarized under application of orthogonal forces. Thus, Flippers have been designed as molecular sensors for compressive forces.

**Figure 1:**
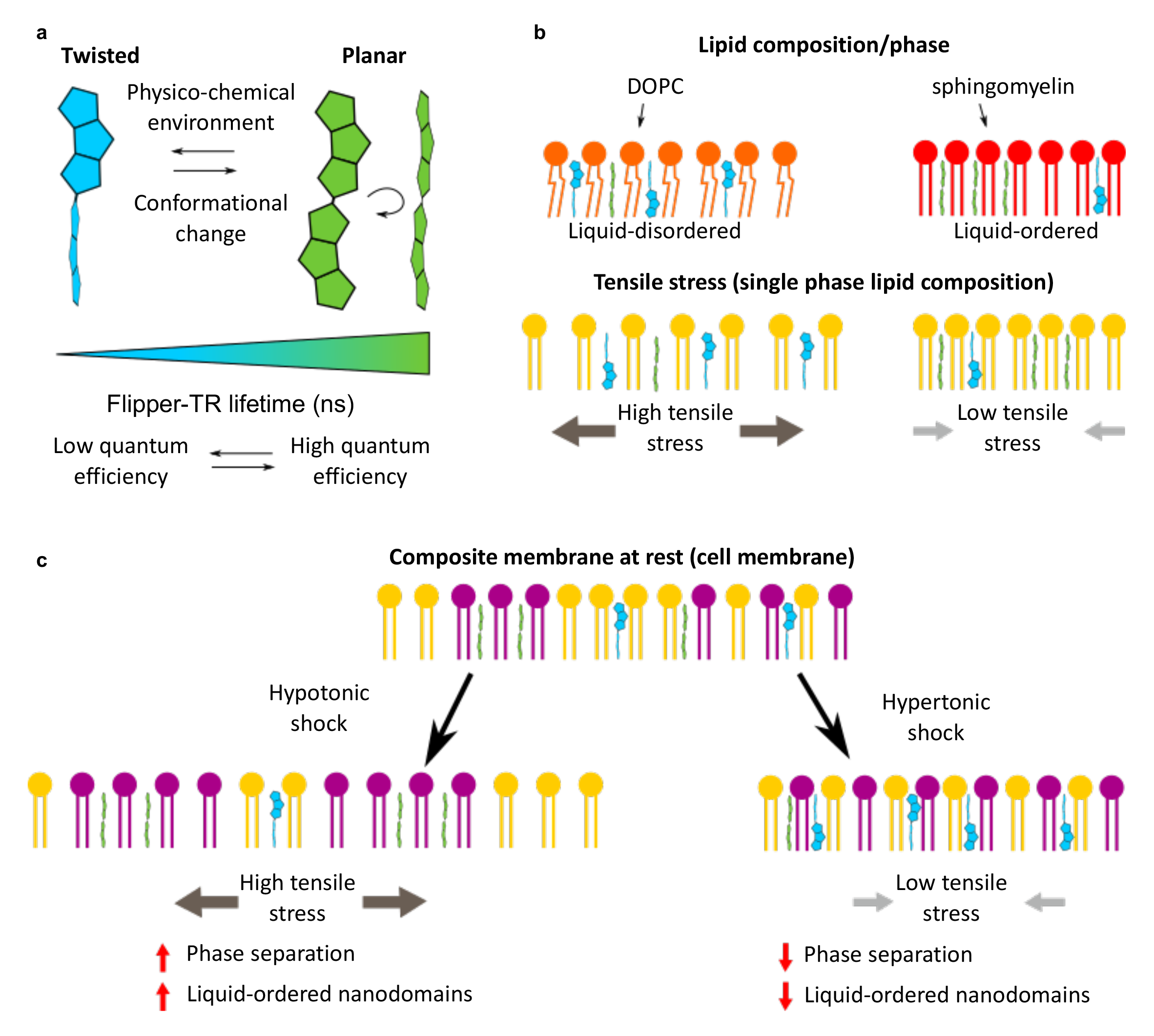
**Principle of Flipper probe** (a) Flipper design is a planarizable push-pull system. Mechanical planarization is responsible for a spectrum of configurations from planar to twisted configuration, which will affect the photo-physics and notably the time of the excited state of the probe (b) Lateral forces within the membrane directly set the twisting and therefore the photo-physics of the Flipper probes. (c) In complex membrane (e.g cell membranes), high tensile stress will trigger phase separation and the formation of highly packed membrane nanodomains reported by an increase of Flippers lifetime.

As the Flipper structure is mostly hydrophobic, the molecule spontaneously inserts in between hydrophobic tails of lipids that constitute cellular membranes (Fig 1b). When inserted, the hydrophobic forces that pack the lipids into a bilayer exert pressure onto the Flipper fluorophore and planarize it. Its planarization depends on the lipid composition, and in short, more ordered membranes exert more force and provide more planarization of Flippers and reversely, disordered membrane, less force and planarization.

The conformation (i.e. how much the Flipper molecule is planarized versus twisted) of the Flipper affects several parameters of the photo-physics of the molecule: in the planarized state, the photon emission efficiency is increased by 10x compared to the most twisted state, and the peak of photon absorption is shifted towards larger wavelength, while the change is minimal for the emission peak [23].

In particular, we noted that the lifetime of Flipper dramatically varied with lipid composition. The lifetime (τ_1_ or the longest decay of a bi-exponential fit) in the twisted state, equivalent to highly disordered lipid membranes, was as small as 2.3 ns, while it could reach values as high as 7 ns in highly ordered membranes, where the molecule is fully planarized. This large range of lifetime values is due to the fact that in the planarized state, fluorescence of the two fluorophores is coupled through an electron transfer from the donor group to the acceptor group[6], which delays emission of the photons, giving longer lifetimes.

The range of lifetime values that this probe could cover was impressively large compared to other probes. Flippers are so sensitive that we imagined that they could report tiny changes of the hydrophobic pressure linked to changes of membrane tension. In cell membranes or in GUVs with phase separating lipid composition, increasing tension results in higher lifetime and decreasing tension results in a shorter lifetime. This is likely due to a tension induced lipid phase separation and membrane reorganization in highly ordered microdomains[24–26] (Fig 1c). These behavior of Flippers in cell membrane have been confirmed in many studies[6,8–10,12,27] and validated by molecular simulation[28]. Flipper is thus complementary to Laurdan, an organic fluorescent maker which allows by measuring its fluorescence anisotropy or its lifetime to detect changes in plasma membrane fluidity[29,30]. Since Flippers lifetime fluctuates with both the lipid composition and membrane tension, changes of the lifetime can be directly associated with a change in tension only in cases where lipid composition is not changing. While this condition is always met *in vitro*, in cells, we usually consider that any variation of lifetime on the second timescale is related to membrane tension, as lipid composition hardly changes over this timescale. For longer timescales (minute, hour), appropriate controls that the lipid composition is not dramatically changing are required.

## II-Staining biological samples with Flipper probes

Being hydrophobic, the probe is in the micellar form in aqueous solutions and is not fluorescent in this state. When the Flipper probe diffuses within lipid membranes, it inserts vertically between lipids and fluoresces. Thus, Flipper probes are "fluorogenic", becoming fluorescent only when inserted in the structure they target. We describe staining procedures, problems and solutions below.

### A-Staining of cultured cells

To label most of the cell lines tested (Fig 2a), the probe, solubilized in DiMethylSulfOxide (DMSO), was normally directly added to the imaging medium at 1 µM final concentration and incubated 15 min at 37°C with cells. MDCK-II cell medium was replaced 1 h before imaging with medium containing 1 mM of Flipper-TR in order to allow the probe to penetrate through the cell monolayer (and, when relevant, the alginate wall). To keep a low concentration of DMSO in the final medium, 1 mM stock solutions of Flipper probes were usually made. In some cell types (MDCK, HeLa Kyoto and MZ), 5 min of Flipper incubation at 37°C might be enough to obtain good staining. However, at high cell density or in 3D cell cultures and tissues, longer incubation - in the order of a few hours - may be necessary. Imaging medium (Fluorobrite with HEPES or CO_2_ supply, or Leibovitz if no CO_2_ supply is available) was usually used and gave a good signal over noise ratio. The use of Foetal Bovine Serum (FBS) during staining and imaging can cause a lower signal, as it contains proteins which can extract hydrophobic molecules from membranes such as Albumin. However, we haven’t observed a dramatic change of labelling efficiency using solutions described above supplemented with FBS. If the staining appears to be low with FBS, we nevertheless recommend testing solutions without it. We did not observe drastic differences on the fluorescence intensity and lifetime of the probe between cells imaged at 37°C and the ones imaged at room temperature. Similarly, CO_2_ adjunction in the imaging chamber or working with a medium supplemented with HEPES does not significantly impact Flipper fluorescence intensity or lifetime.

**Figure 2:**
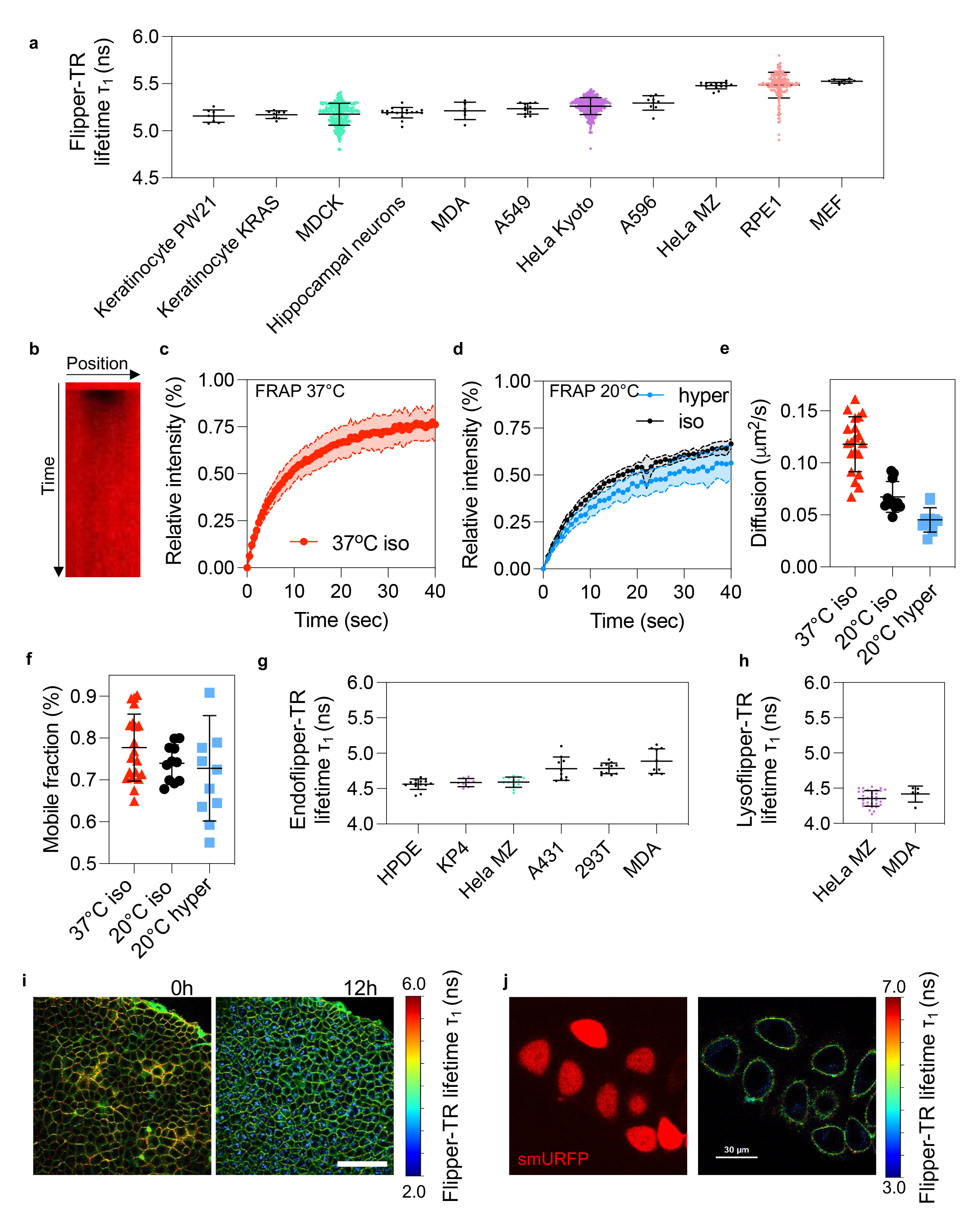
**General behavior of Flipper probes** (a) Graph showing the Flipper-TR lifetime (τ1) extracted from different cell types (b) Kymograph showing the recovery of Flipper-TR signal after bleaching at the plasma membrane (FRAP experiment) (c) Graph showing FRAP curves of Flipper-TR at the plasma membrane at 37°C on HeLa cells in isotonic medium (n = 18) (d) Graph showing FRAP curves of Flipper-TR at the plasma membrane at 22°C on HeLa cells in isotonic (320 mOsm) medium (n = 11) versus hypertonic (800 mOsm) medium (n = 10) (e) Graph showing diffusion time coefficients (μm^2^/sec) extracted from graph (d) data (f) Graph showing the mobile fractions (%) extracted from graph (d) data (g) Graph showing the Lyso Flipper lifetime (τ1) extracted from different cell types (n>5 fields of view with at least 5 cells per field for each condition) (h) Graph showing the Endo Flipper lifetime (τ1) extracted from different cell types (n>5 field of view with at least 5 cells per field for each condition) (i) FLIM images showing MDCK tissue stained with Flipper-TR at different timepoints. (j) Flipper-TR FLIM and confocal image of HeLa cells labelled with Flipper-TR lifetime and smURFP fluorescence

Importantly, the cell labelling occurs through a dynamic equilibrium of the probe with the micellar pool in the cell culture medium. Thus, a constant exchange of Flipper-TR between the plasma membrane and the medium occurs. This exchange is rapid for the Flipper-TR because the medium is directly in contact with the plasma membrane labelled by the probe.

Using FRAP without Flipper-TR in the medium, we detected a faster diffusion coefficient D at 37°C, D= 0.12 +/- 0.04 μm^2^/sec (mobile fraction of 78% +/- 5) (Fig 2b, c) than at 20°C, D=0.07 +/- 0.02 μm^2^/sec (Fig 2d, e). As expected, a decrease of plasma membrane tension induced by hypertonic treatment reduces the Flipper-TR diffusion coefficient at room temperature from 0.07 +/- 0.02 μm^2^/sec to 0.05 +/- 0.01 μm^2^/sec (Fig 2e) without significantly impacting the mobile fraction (Fig 2f).

We recommend, if possible, to keep a constant Flipper-TR concentration along the experiment. Concerning Lyso Flipper-TR, the molecule diffuses freely across the plasma membrane and quickly concentrates in the membrane of late endosomes and lysosomes (in a few minutes) after headgroup protonation in this acidic environment. Consequently, for short timescale experiments (less than 2h) it is not an issue to leave cells in Flipper medium containing Lyso Flipper-TR. However, loss of the acidity of endosomal compartments after treatments or change in the pH of the medium (due for example to phototoxicity) may trigger loss of the Lyso Flipper-TR staining. The best labelling was obtained by incubating Lyso Flipper-TR at 1 µM in Fluorobrite medium without serum for 20 minutes at 37°C prior to imaging.

Because of the dynamic exchange of the probe between the membrane and the medium, washing Flipper labelled cells with medium containing FBS or BSA will decrease Flipper-TR staining. Therefore, if washing is absolutely needed, it is recommended to wash with medium without FBS, and/or keeping the concentration of the Flipper constant in the washes, which will limit the decrease of the membrane signal.

Reaching the dynamic equilibrium takes about 5-10 minutes, thus the photon count depends on the time of incubation of cells with the probe, until it reaches a plateau value that depends on your system. Incubations that are too long may lead to Flipper-TR endocytosis and endosomal labelling. The Flipper-TR lifetimes within endosomal compartments are notably lower than in the plasma membrane (Fig 2g, h), and we have never observed Flipper-TR labelling expanding further away than from the endosomes (probably even the early endosomes). Also, the Flipper-TR endosomal labelling appeared with various times depending on the cell types and experimental conditions. In MDCK, we observed that endocytosis occurred after 12 hours (Fig 2i).

For long-term imaging experiments (several hours) with sufficient time-lapse between images, we recommend performing sample-labelling for each time point: the sample is incubated with Flipper-TR, imaged, and then the probe is washed away using medium with BSA or FBS. Each timepoint requires re-incubation of the probe to prevent Flipper-TR endocytosis.

### B-Co-staining with Flipper-TR

In order to specifically analyse the tension or lipid composition of a given organelle or during a biological process, it is possible to use other fluorescent probes or stable cell lines in conjunction with Flipper-TR. However, to avoid any fluorescence overlap with the probe, which could affect lifetime measurements, blue or far-red fluorescent probes should be used (e.g: Organic dyes like Alexa-405 or Alexa-647 or Blue fluorescent protein, E2-Crimson or smURFP for fusion proteins, see Fig 2j). Indeed, due to the configuration of the probe (two heads), Flipper-TR has a large spectrum of emission and a large spectrum of absorption. The excitation peak of Flipper is at 485 nm and fluorescence emission is collected using a band pass 600/50 emission filter. Therefore, for example in the case of an organelle labelled with Alexa-405, excitation of this fluorophore (peak at 400 nm) will only weakly excite Flipper-TR, limiting the phototoxicity while the emission of Alexa-405 (400-530 nm) will not go through the FLIM detector and will not interfere with Flipper-TR lifetime measurement.

### C-List of cells and organisms from which we obtained successful staining and imaging using Flipper probes

#### List of cell types used with Flipper-TR

HeLa MZ, HeLa Kyoto, MDCK, keratinocyte PW21, keratinocyte KRAS, hippocampal neurons, MDA, A549, A596, RPE1, MEF with lifetime value (tau1) ranging from 5 ns to 5.7 ns (Fig 2a).

#### List of cell types used with Lyso-Flipper-TR

HPDE, KP4, Hela MZ, A431, 293T, MDA with lifetime value (τ_1_) ranging from 4.4 ns to 5.1 ns (Fig 2g).

#### List of cell types used with Endo Flipper-TR[6,31]

HeLa MZ, MDA with lifetime value (τ_1_) ranging from 4.1 ns to 4.7 ns (Fig 2h).

#### List of organisms stained with Flipper-TR

alginate capsules containing cells, PDMS tubes with attached cells on the inner surface, Arabidopsis thaliana leaf, Arabidopsis thaliana plant root, Bacillus Subtilis (Fig 3i), Xenopus explant (Fig 3j), 3D gastruloids of mESC (Fig 3k) or mouse embryos (Fig 3l and ref[32]).

**Figure 3:**
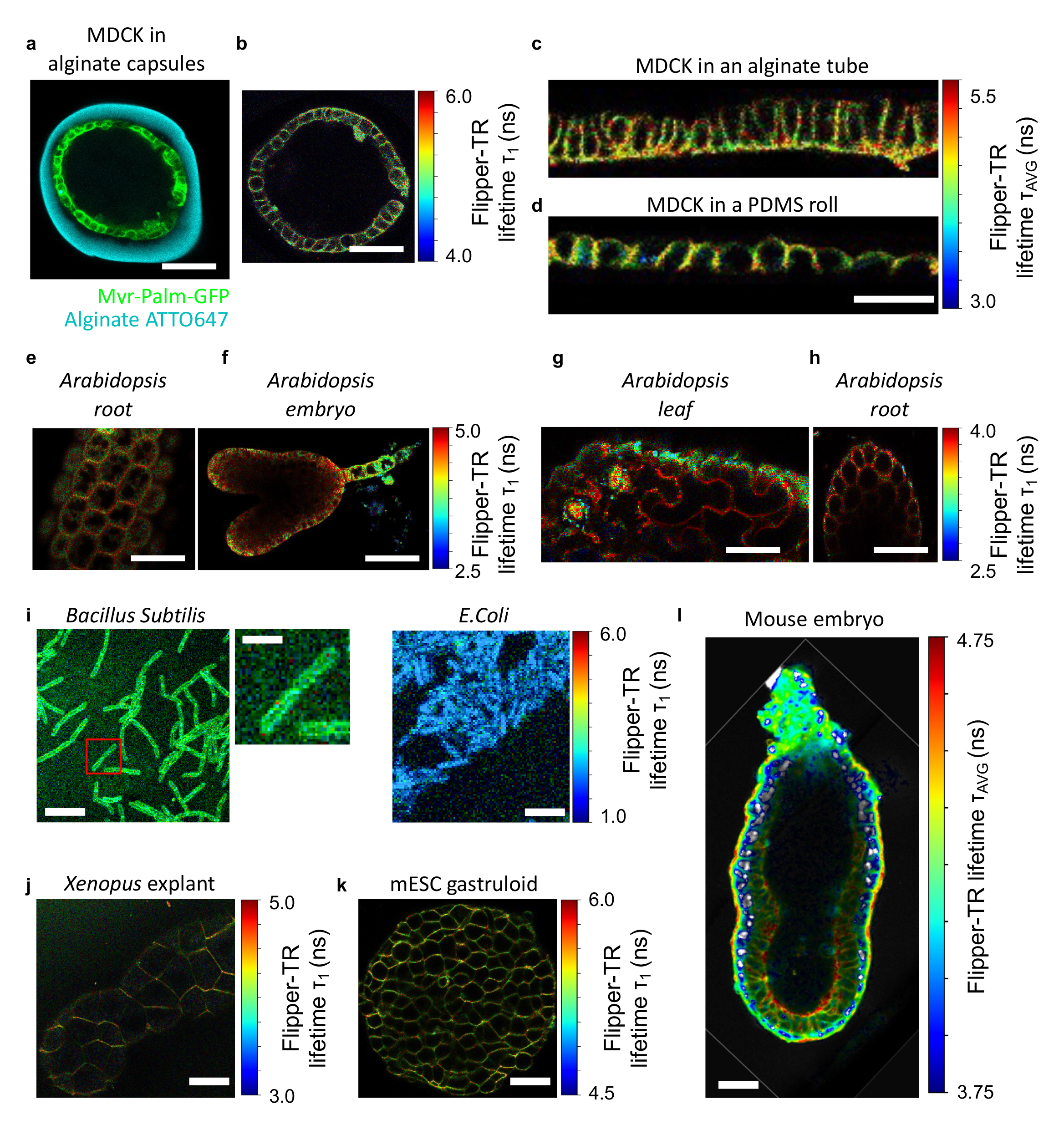
**Flipper probes staining in various model and organisms** (a) Representative fluorescence image of a MDCK monolayer in an alginate capsule (Myr-Palm-GFP stain the MDCK cells and ATTO647 stain the alginate capsule in blue). Scale bar is 50 μm. (b) Representative FLIM image of an MDCK monolayer stained with Flipper-TR in an alginate capsule. Scale bar is 50 μm. (c) Representative FLIM bottom view of an MDCK monolayer stained with Flipper-TR in a alginate tube. (d) Representative FLIM side view of an MDCK monolayer stained with Flipper-TR in a PDMS roll. Scale bar is 20 μm. (e) Representative FLIM image of Arabidopsis root stained with Flipper-TR. Scale bar is 20 μm. (f) Representative FLIM image of the entire Arabidopsis embryo stained with Flipper-TR. Scale bar is 40 μm. (g) Representative FLIM image of Arabidopsis leaf stained with Flipper-TR. Scale bar is 20 μm. (h) Representative FLIM image of Arabidopsis root stained with Flipper-TR. Scale bar is 40 μm. (i) Representative FLIM image of Gram-positive (B. subtilis) stained with Flipper-TR and representative FLIM image of Gram-negative (E. coli) stained with Flipper-TR showing a non-specific signal. Scale bar is 10 μm. (j) Representative FLIM image of Xenopus explant stained with Flipper-TR. Scale bar is 100 μm. (k) Representative FLIM image of mESC gastruloid stained with Flipper-TR. Scale bar is 80 μm. (l) Representative FLIM image in E6.5 mouse embryos stained with Flipper-TR. Scale bar is 20 μm.

List of organisms where Flipper-TR staining does not work: E.Coli (gramm – cells) (Fig 3i, right panel). E.Coli showed only background Flipper-TR staining when compared to Bacillus Subtilis, suggesting that the presence of a cell wall in gram negative bacteria could prevent Flipper-TR staining. Different attempts were made by several groups to stain zebrafish but they were unsuccessful so far.

### D-Different probes for different staining

Various derivatives of Flipper were developed to target the endoplasmic reticulum (ER Flipper-TR), the mitochondria (Mito Flipper-TR), the lysosome/late endosome (Lyso Flipper-TR)[22] and more recently the early endosome (Endo Flipper-TR)[31] as well as a version of the probe for single-molecule super-resolution imaging of membrane tension (SR-Flipper[27,33]). The structures can be found in[6]. The ER Flipper-TR selectively labels the membranes of endoplasmic reticulum via a pentafluorophenyl group which reacts with cysteine of proteins present on ER outer surface. The average lifetime of ER Flipper-TR is lower (3.5 ns in HeLa cells) than Flipper-TR (4.5 ns in HeLa cells) in various cell lines. The Mito Flipper-TR (Spirochrome)[6,22]; selectively labels the membranes of mitochondria via the interaction between the hydrophobic triphenylphosphonium cation with the negatively charged surface and the inside negative potential of the mitochondrial membrane. As observed for ER Flipper-TR, the average lifetime of Mito Flipper-TR is rather low (around 3.2 ns in HeLa cells).

Lyso Flipper-TR and Endo Flipper TR contain a morpholine headgroup of higher pKa in Endo Flipper-TR compared to Lyso Flipper-TR. This morpholine group is protonated in the acidic environment of endosomes and retained in its membrane. The average lifetime of Lyso Flipper-TR is around 4 ns in HeLa cells. Endo and Lyso Flipper-TR are showing very weak phototoxicity.

SR-Flipper[33] is behaving as Flipper-TR (labelling plasma membrane in cells) and is reversibly switching from bright-state ketones to dark-state hydrates, hemiacetals, and hemithioacetals both in twisted and planarized state. It is therefore possible to use it for single-molecule localization microscopy and to resolve membranes well below the diffraction limit. The lifetime (τ_1_) of SR-Flipper is usually slightly lower (∼10%) than the lifetime of Flipper-TR. HaloFlipper is based on the combination of the Flipper and a Halo tag to label any membrane of interest by targeting membrane specific protein labelled with a halo tag. Halo PhotoFlipper targets the nuclear membrane and the inner plasma membrane using a photocleavable domain[34].

## III-Acquisition

Flipper probes have long average fluorescence lifetimes, up to 7 ns depending on membrane composition, which means that photons with lifetime up to 50 ns can be detected. Consequently, FLIM systems with sampling frequencies of 20 MHz are the best devices to use to detect all the photons allowing for most accurate TCSPC fits. We found however that the dye can also be efficiently excited using a two-photon mechanism (excitation around 880 nm – 950 nm, Fig S1a, b). Unfortunately, most two-photon microscope set-ups are only equipped with a fixed repetition rate laser pulsing at 80 MHz. Increasing the pulse rate leads to artificially shortened lifetimes in GUVs (excited with one-photon at 488 nm) (FigS1c, d, e). While the lifetime of Flipper-TR in GUVs still depends on the membrane compositions, the dynamic range is reduced at 80 MHz compare to 20 MHz (FigS1c,d,e). The use of Flipper-TR in conjunction with the possibility of two-photon excitation is a promising feature for *in vivo* studies and deep tissue imaging of tension.

On PicoQuant systems the pulsed laser frequency can be set between 80 MHz and 31.25 kHz, but on LEICA systems the laser frequency is by default 80 MHz which will not allow to measure lifetimes higher than 12.5 ns which will represent a substantial amount of photons for a fluorophore with an average lifetime a 4 ns for instance. Therefore, the sampling frequency has to be reduced in LEICA systems, using the pulse-picker device (illuminates one out of 2 or more laser pulses), otherwise the lifetime will be underestimated.

**Figure S1:**
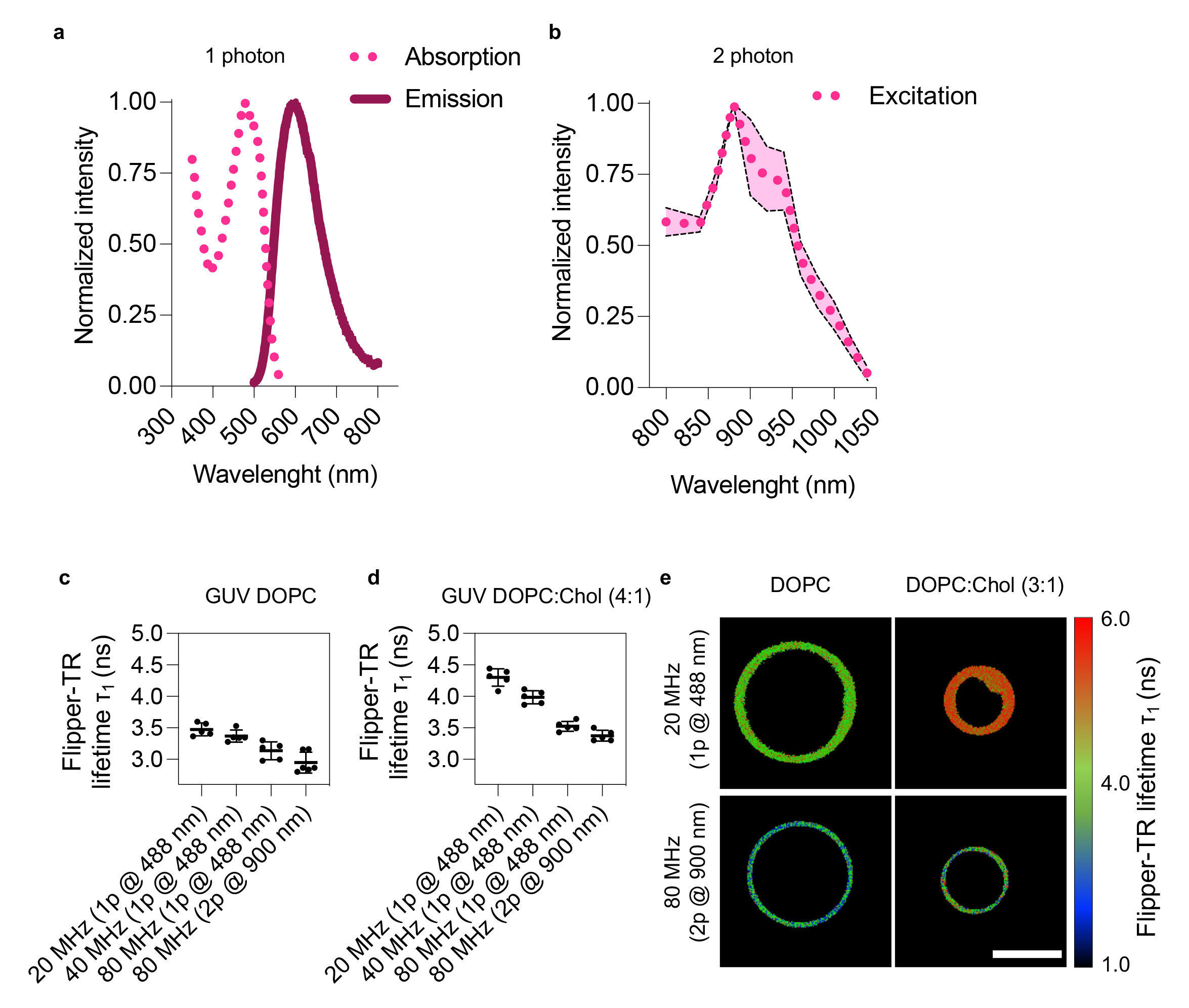
**FLIM of Flipper-TR using two-photon microscopy** (a) Absorption and excitation spectrum of the Flipper-TR. (b) Two-photon excitation spectra of Flipper-TR, acquired in 48h gastruloids. Five gastruloids were imaged in total at each wavelength. The acquired spectra were normalized to the maximum intensity value. Dots represented the average value, scattered lines the minimum/ maximum values of the 5 recorded spectra. (c) and (d): Lifetime of Flipper-TR in GUVs of different compositions DOPC (c) and 1:4 DOPC:Cholesterol (d) at different laser repetition rates and under two-photon excitation. Every dot represents fitted lifetime for 1 GUV, error bars are standard deviation and line is mean. (e) Fluorescence lifetime images of GUVs from (c,d) under typical excitation conditions (20 MHz, 488 nm, top panel) and under two-photon excitation (80 MHz, 900 nm, bottom panel). Longer lifetime component is shown (shorter lifetime component fixed). Image size is 36.4*36.4 µm^2^

Because of the bi-exponential nature of the Flipper decay curve, it is important to record a sufficient number of photons without saturating the detector. We recommend recording a minimum intensity peak of 10^4^ photons per field of view and with at least 200 photons for the brightest pixel.

Another important point is that the phototoxicity during acquisition is lower when the laser pulse frequency is decreased. With classical Time Correlated Single Photon Counting (TCSPC) it is very important to have a count rate not exceeding 1 to 5% of the excitation rate (for 20MHz pulse, average detector count rate should not exceed 1 MHz) in order to maintain a low probability of registering more than one photon per excitation cycle (with the dead time, the system would detect the first photon and miss the second one, called “pile up” effect).

Other classic confocal microscopy parameters that participate in light acquisition (pinhole aperture, laser power, pixel binning, scanning speed, size of the ROI, line/frame summation) may be used to optimize photon collection. Several means can be used to reduce the phototoxicity:

- Using a new generation of TCSPC device with reduced dead time (rapidFLIM from Picoquant, dead time of less than 650 ps compare to 80 ns in normal FLIM) that allows to detect more than one photon per excitation cycle. It also provides a better temporal resolution (see below) and allows to work with higher intensities. Overall, with this new kind of TCSPC much higher detector count rates can be processed which result in a faster acquisition. This system also allows to acquire more than one photon per laser pulse without the pile up effect.
- Repeating acquisition and summing-up photons from several images will allow to obtain sufficient counts of photons (minimum peak of 10^4^ events) for a reliable fit. For standard cells in culture, we used the following conditions. For example, using Flipper-TR in HeLa Kyoto, we could sum up to 7 frames in a 256×256 pixel image to reach 10^5^ photons peak in case of an entire field of view which was necessary for fitting procedure and lifetime extraction. Cell type and signal intensity determine the optimal acquisition parameters. It is necessary to find a reasonable balance between photon number and phototoxicity and we created a troubleshooting table based on our experience (Fig 4).

**Figure 4:**
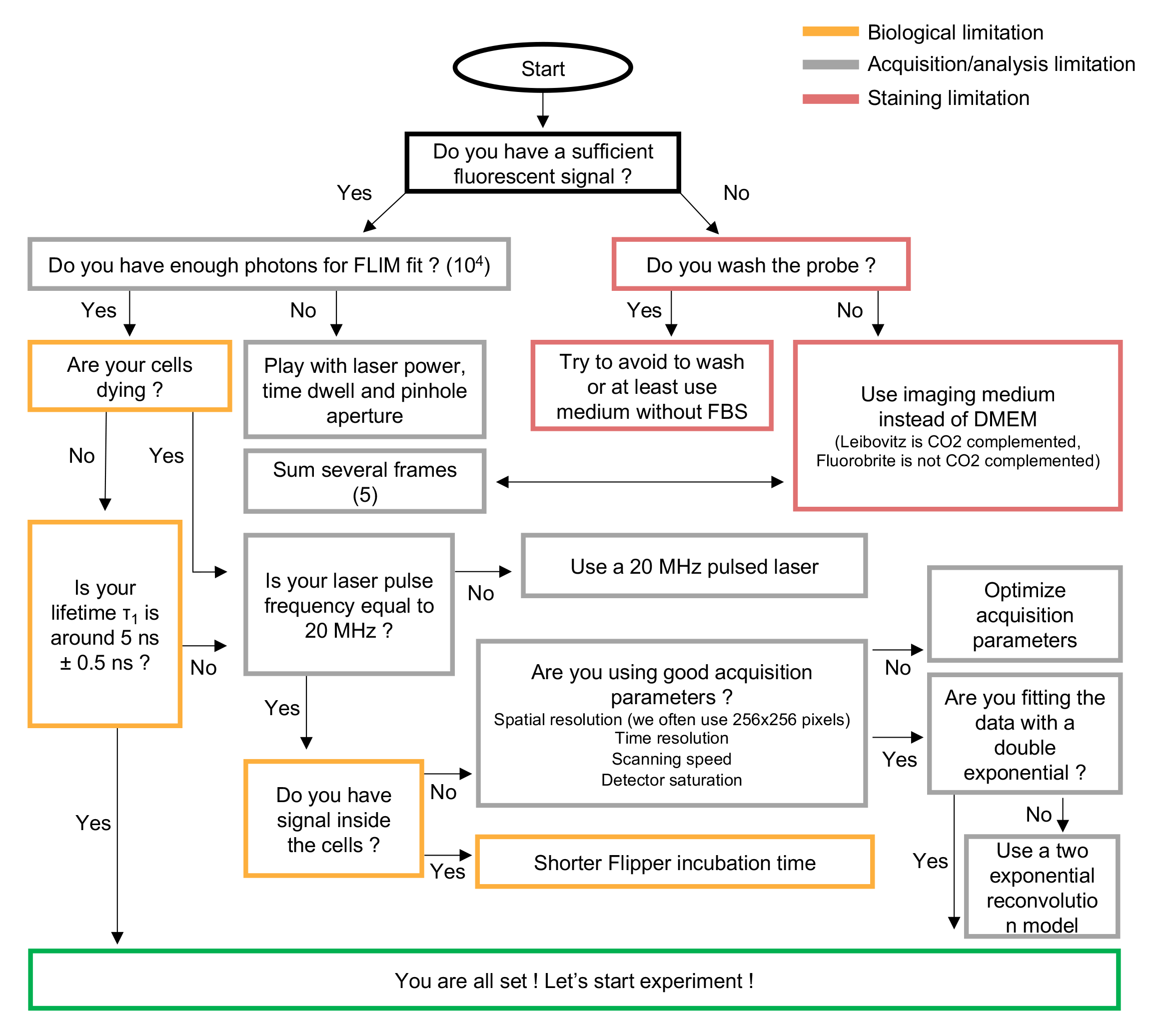
**Chart to support users with troubleshooting Flipper probes experiments.**

### Timelapse of Flipper-TR

Since the Flipper-TR probe is in dynamic exchange with the environment, measuring lifetime over time has its own challenges. If medium is added (osmotic shocks or drug addition), we advise to keep Flipper-TR concentration constant. If medium is flowed (to test the effect of shear stress for example) it is essential to keep the Flipper-TR concentration constant. Because lifetime values are broad due to inherent biological variability and impacted by cell density or cell type, we suggest assessing both lifetime variation and absolute lifetime values.

### Combining Flipper probes with drug treatment

To measure the effect of a drug treatment on cells, we tested two strategies. (1) Imaging the cells, adding the drug in the medium of a dish or replacing the medium with the one that contains the drug using a microfluidic device and following the effect of the drug over time on the same cells. This approach allows to eliminate the biological variability by measuring the tension of the same cells before and after the treatment. (2) Imaging the control condition and the treated condition independently (preferably acquire a large amount of cells to increase statistics as it will be unpaired measurements, typically at least 15 fields of view per dish and 3 technical replicates). The treated cells must be imaged in the same conditions and on the same day. Because steady state lifetime value can be variable (it depends on cell density), the control condition must be repeated along every drug condition with the same seeding parameters. We suggest following strategy (1) whenever possible. Classical solvents (DMSO) or antibiotics used in inducible systems (doxycyclin) change the property of the membrane and therefore affect the Flipper-TR lifetime. Indeed, DMSO increases solution hypertonicity while doxycyclin affects cholesterol amounts.

### Acquiring lifetime in a z-stack (such as in polarized tissue)

We observed a weak staining in very dense and polarized tissue independently of the incubation time (from 5 min to 2h). Tissues are not perfectly flat leading to smaller collection of photons at the apical side, which might lead to inhomogeneous signal and thus poorly reproducible results. To compare lifetime of the Flipper-TR along Z-axis, samples and conditions, we suggest defining the basal or mid-plane as the plane with the largest number of photons. Due to the inherent difficulty to measure the lifetime of single cells, we advise to compare planes containing several cells.

## III-Analysis

### A-Tools to analyse the data

Several companies have made commercial packages for FLIM analysis, but these are closed source tools that are not transparent in their analyses and typically only support their own file formats. Open source and user friendly tools have been and are being developed[35,36].

### B-IRF (instrument response function)

In order to fit the signal, the contribution of instrumentation must be isolated. The overall timing precision of a complete TCSPC system is its Instrument Response Function (IRF). The best way to measure the IRF of the system is to use a fluorophore with similar fluorescence properties as the Flipper-TR but with a very short lifetime. The IRF can be measured using fluorescein solution quenched with potassium iodide before each experiment or can be calculated (deducted) from the rising edge of the TCSPC histogram. The important point is to use the same IRF throughout the experiment. Software used to analyse FLIM images usually recalculate the IRF for each file by default.

### C-Photon count fitting

Two different methods to fit the exponential decay exist: exponential reconvolution and exponential tailfit. A tailfit can be used when the lifetimes are significantly longer than the IRF. In general, a reconvolution fit is preferable, because the complete decay is fitted, while the start of the fitting range is slightly arbitrary for a tailfit. Depending on the format of the data and the available tools, choosing one or the other should not impact measurements too much if enough photons are acquired. Tens of thousands of photons per pixel are required to accurately fit a bi-exponential decay [37]. In general, lifetime histograms of Flippers are not well fitted by a mono-exponential model, so it is standard to fit the data with double exponentials. By fitting with a bi-exponential decay, two lifetimes will be extracted: τ_1_ and τ_2_. τ_AVG_ is an average between the two rates τ_1_ and τ_2_ weighted by the number of photons fitted by each parameter. τ_1_ (the longest component, usually ∼80% of photons with a lifetime around 5.5ns for Flipper-TR at the plasma membrane in HeLa) usually represent much higher photon counts than τ_2_ (usually ∼20% of photons with a lifetime around 1ns for Flipper-TR at the plasma membrane in HeLa). Both τ_1_ or τ_AVG_ will directly report the mechanical property (while the fluctuation of τ_2_ will be less pronounced) of the probe so you should anyway take either τ_AVG_ or the longest τ (τ_1_ on picoquant system) value to analyse tension fluctuations. In experiments analyzing the dynamics of Flipper, if the tendency of τ_1_ and τ_AVG_ are different, probably because of a lack of photons, the fit and image acquisition must be optimized. In some cases, fit free approaches can be used as well. The fast lifetime (often defined as the arithmetic mean on the photon arrival times per pixel) can be used and shows similar values to the fitted average lifetime τ_AVG_.

The exponential fitting can be done on a different set of pixels. The choice depends on the quality of your signal and the sensitivity of the effect you are looking at. (1) Whole photons analysis is the simplest and necessitate little requirement of input from the experimentalist, therefore increasing the reproducibility of the work. (2) It is possible in order to remove the background to apply a threshold on pixel intensity and only fit the corresponding pixels. It is however invalid to apply a threshold based on the lifetime values to extract lifetime. (2’) Applying a threshold on the absolute lifetime value can be valid to create masks and separate objects with different Flipper lifetimes. For example, the lifetime of the plasma membrane is much higher than the lifetime of endosomal membrane, so a single FLIM image could be sufficient to discriminate the two objects. (3) Similar to (2), it is possible to overlap a mask generated from another channel to only select pixels of interest (Fig 5a-b).

**Figure 5:**
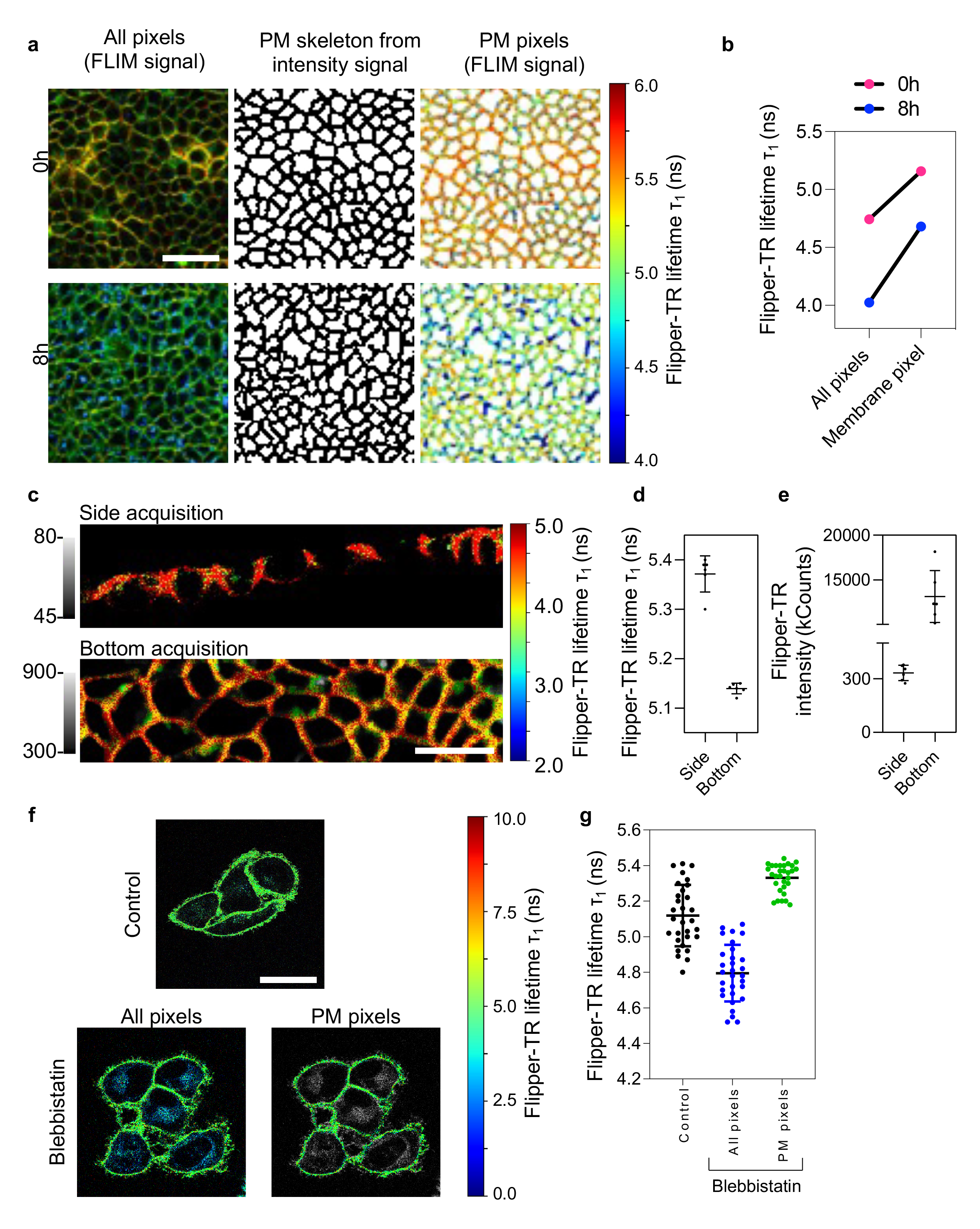
**How to properly extract the Flipper-TR lifetime from FLIM image.** (a) Representative FLIM images of MDCK tissue stained with Flipper-TR (top row) with and without endosomal staining (respectively top panel and bottom panel after 8h of incubation) due to longer cell incubation with Flipper-TR. Flipper-TR lifetime of pixels corresponding only to plasma membrane (PM) extracted via the segmentation of the PM (middle column) based on Flipper-TR fluorescence intensity. (b) Graph showing the Flipper-TR lifetime (τ_1_, ns) extracted from all the pixels of the image versus only the pixels corresponding to the plasma membrane. (c) Representative FLIM images of MDCK tissue in PDMS roll stained with Flipper-TR acquired from the side (top row) or from the bottom (bottom row). (d) Graph showing the Flipper-TR lifetime (τ_1_, ns) extracted from the pixels of the side acquisition image versus from the bottom acquisition image. (e) Graph showing the number of photons extracted from the pixels of the side acquisition image versus from the bottom acquisition image. (f) Representative FLIM images of HeLa cells incubated with DMSO (top) or with blebbistatin (bottom). Bottom images show two configurations: all the pixels are selected (left) or only pixels corresponding to PM were manually selected (right). Scale bar is 40 μm. (g) Graph showing Flipper-TR lifetime quantification corresponding to (c). Lifetime (τ_1_, ns) was extracted from all the pixels of the image versus only the pixels corresponding to the plasma membrane (n=30 for both control and blebbistatin treated cells)

**Figure S2:**
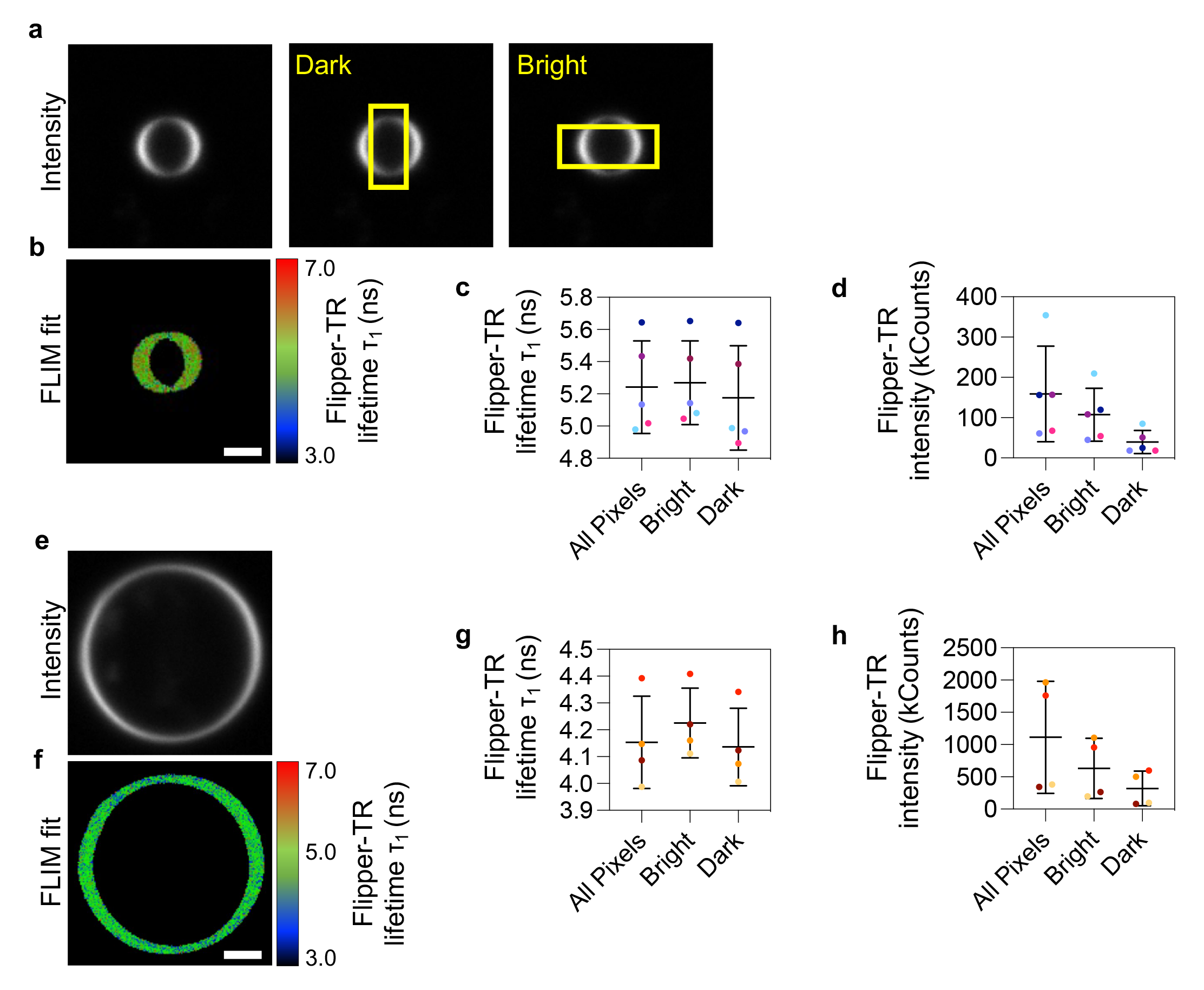
**Effect of linearly polarised excitation light on Flipper-TR lifetime measurements in GPMVs and GUVs.** (a) Intensity and (b) Lifetime images of cell-derived vesicles. Photoselection of the fixed excitation dipole of the Flipper-TR probe (using a linearly polarized excitation laser) causes bright rings with higher intensity. Graphs show (c) Lifetime and (d) intensity analysis of bright vs dark regions. Every dot represents the lifetime fit across one vesicle. The line represents the mean and error bars are standard deviation. (e) Intensity and (f) lifetime and image of artificial vesicle composed of DOPC and Cholesterol (4:1). Image size is 36.4*36.4 µm^2^. The graphs represent (g) lifetime and (h) intensity analysis of bright vs dark regions. Every dot represents the lifetime fit across one vesicle. The line represents the mean and error bars are standard deviation.

Depending on the orientation of the cells in the beam, the number of photons can change (Fig S2). A side acquisition will give a lower number of photons than an acquisition from the bottom, leading to a worse fit (Fig 5c-e). As can be noticed from the intensity images and quantification (Fig S2 a, e, d, h), Flipper-TR undergoes photoselection when excited with linearly polarized light. This effect can be misleading for probes like Laurdan that use fluorescence intensity to report on membrane properties[38]. While the photoselection during excitation can lead to bright edges of vesicles or membranes aligning or misaligning with the polarization, fluorescence lifetime measurements and fitting (given enough photons are acquired) is not significantly different between bright and dark regions (FigS2 c, g) and can robustly report on the membrane properties. Note that the effect is stronger for membranes with higher viscosity (larger cholesterol content). However, the photoselection effect can be misleading when using intensity-weighted lifetime representations (as in FastFLIM).

### D-Image segmentation to extract Flipper lifetime

Here, we will develop two examples to illustrate the importance of extracting the lifetime properly. Fluorescence intensity images of a membrane probe (CellMask Deep Red for instance) can be used to generate a mask that permits to extract lifetime of all the pixels present in the mask (Fig 5a). Even if the lifetime of the internalized probe is lower than the lifetime of the probe at the plasma membrane, applying a threshold based on lifetime value will introduce a bias in the analysis of the overall lifetime. Considering only pixels corresponding to the plasma membrane instead of all the pixels of the image, which include signal coming from endosomes, gives a difference of 0.5 ns on average (Fig 5b).

The second example is illustrated by measuring the impact of blebbistatin (myosin inhibitor reducing cell contractility) on plasma membrane tension reported by Flipper-TR lifetime (Fig 5f). If all the pixels of the images (control versus blebbistatin-treated cells) are included for the fitting, a decrease of lifetime of 0.3 ns is observed (Fig 5g). Each dots represent a field of view containing several cells, the fitting was performed by fitting all the pixels. However, it appears that on images of Blebbistatin-treated cells, despite an identical Flipper-TR incubation time, many intracellular membranes with a low lifetime around 2.5 ns are visible. These pixels likely contribute to the overall lower lifetime measured for Blebbistatin-treated cells. By contrast, if only plasma membrane lifetime is measured, by selecting the plasma membrane pixels, the average lifetime of Blebbistatin-treated cells is 0.1 ns higher than the average lifetime of control cells. Directly selecting the plasma membrane pixels also excludes the background, which could affect the analysis.

### E-Comparing different areas within the same image

The goal of the experiments might be to compare regions within the same image. As explained above, a minimum number of photons will be necessary to obtain a decent fit and to extract the lifetime properly. Therefore, we strongly suggest checking that the photon count is similar between different area analyzed.

## IV-Results

### A-Different lifetime along cell height

By imaging Flipper-TR over z-stacks of MDCK polarized tissues, we observed a lower lifetime (4.6 ns) at the basal plane of the cell compared to the apical plane (5.4 ns), as illustrated in Fig 6a. Also, large variability exists between position in the dish: the basal plane lifetime values vary from 4.2 ns to 5.1 ns, while the apical plane values vary between 4.3 ns and 5.6 ns. However, the lifetime in the apical plan is reproducibly higher than in the basal plan (Fig 6b) which could be explained by the known differences in lipid compositions. Therefore, single plane imaging of cells to compare, for example, different treatments, must be performed at a similar height (Fig 6c), and sufficient statistics to account for large biological differences between single cells are required.

**Figure 6:**
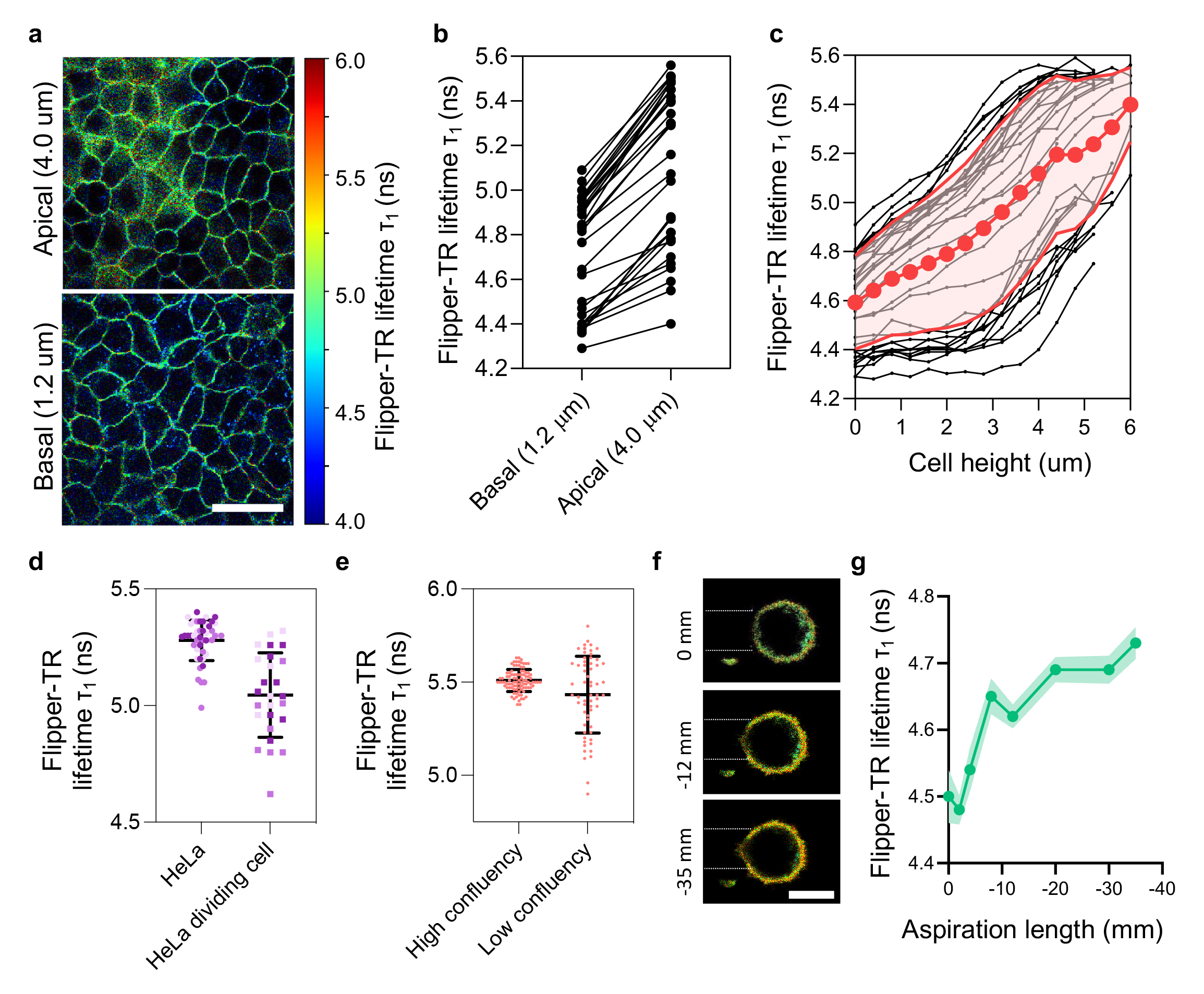
**Flipper-TR lifetime is affected by several parameters** (a) Representative FLIM images of MDCK tissue stained with Flipper-TR in the apical plan (top row) ant the basal plan (bottom row). Scale bar is 40 μm. (b) Graph showing the quantification of Flipper-TR lifetime (ns) in the apical plane (4.0 micron from the bottom) and the basal plane (1.2 micron from the bottom). The bottom was determined as the plane with the highest number of photons (n = 45). (c) Graph showing the quantification of Flipper-TR lifetime (τ_1_, ns) from the bottom plane to the apical plane by steps of 0.4 micron. Each curve represents the lifetime of an entire field of view (n = 45). (d) Graph showing the quantification of Flipper-TR lifetime (τ_1_, ns) in mitotic and non-mitotic HeLa cells (e) Graph showing the quantification of Flipper-TR lifetime (τ1, ns) in confluent and non-confluent RPE1 cells (f) Representative Flipper-TR FLIM images of HeLa cell aspirated in a micropipette at different aspiration pressure. Scale bar: 10 μm (g) Graph showing the quantification corresponding to (c) of Flipper-TR lifetime (τ_1_, ns) depending on the aspiration pressure applied to the cell.

### B-Mitotic cells have a more disordered plasma membrane

We analyzed the Flipper-TR lifetime in mitotic cells which are known to have a higher volume, higher cortical tension and a very different shape than adherent cells[39–41]. Mitotic cells were identified by eye based on their rounded shape and their limited attached surface to the glass bottom. In contrast to the literature, mitotic cells have a 0.35 ns lower lifetime (Fig 6d) compared to adherent cells[40,41], while one might have expected an increase of Flipper-TR lifetime. This lower lifetime could reflect a change of lipid composition during cell division[42], itself causing increased membrane fluidity in mitotic cells[43].

### C-Flipper-TR variability depends on cell confluency

We analyzed the Flipper-TR lifetime depending on cell confluency. Using RPE1 cells, we observed that confluent cells have a Flipper-TR lifetime centered around 5.5 +/- 0.1 ns while non-confluent cells lifetime is centered around 5.4 +/- 0.2 ns (Fig. 6e). Therefore, in case of experiments performed using cells at low confluency, larger statistics are required to account for large biological differences between single cells.

### D- Distinguishing the contributions of lipid composition and tension to the changes of lifetime

Previous work from the lab[8] showed that mechanical increase of Giant Unilamellar Vesicles (GUVs) membrane tension induced by micropipette aspiration (high tension) leads to an increase of 0.2 ns of Flipper-TR lifetime. Using the same setup to aspirate HeLa cells in suspension (Fig 6f), an increase of 0.2 ns was also observed (Fig 6g), confirming that in both GUVs and cells Flipper-TR lifetime measurements can directly report a mechanically induced change of membrane tension.

Although being slightly lower than Flipper (steady state lifetime of 5 ns vs 5.8 ns for SM/Chol GUV), the lifetime of super-resolution (SR)-Flipper also reports changes in membrane tension of GUVs (Fig 7a-b) as well as change in lipid packing (composition). Indeed, SR-Flipper-TR lifetime of SM/Chol (70/30) GUV is centred around 5 ns while the lifetime of DOPC/Chol (70/30) GUV is centred around 4.2 ns (Fig 7b). With both lipid compositions, SR-Flipper lifetime decreases upon membrane tension decrease induced by hypertonic shocks proving that Flipper probes lifetime report both lipid composition and membrane tension (Fig 7a-e). In contrast, the lifetime of pure DOPC GUVs is not changing after hypertonic shock which is in good agreement with an absence of phase separation (Fig 7a,f).

**Figure 7:**
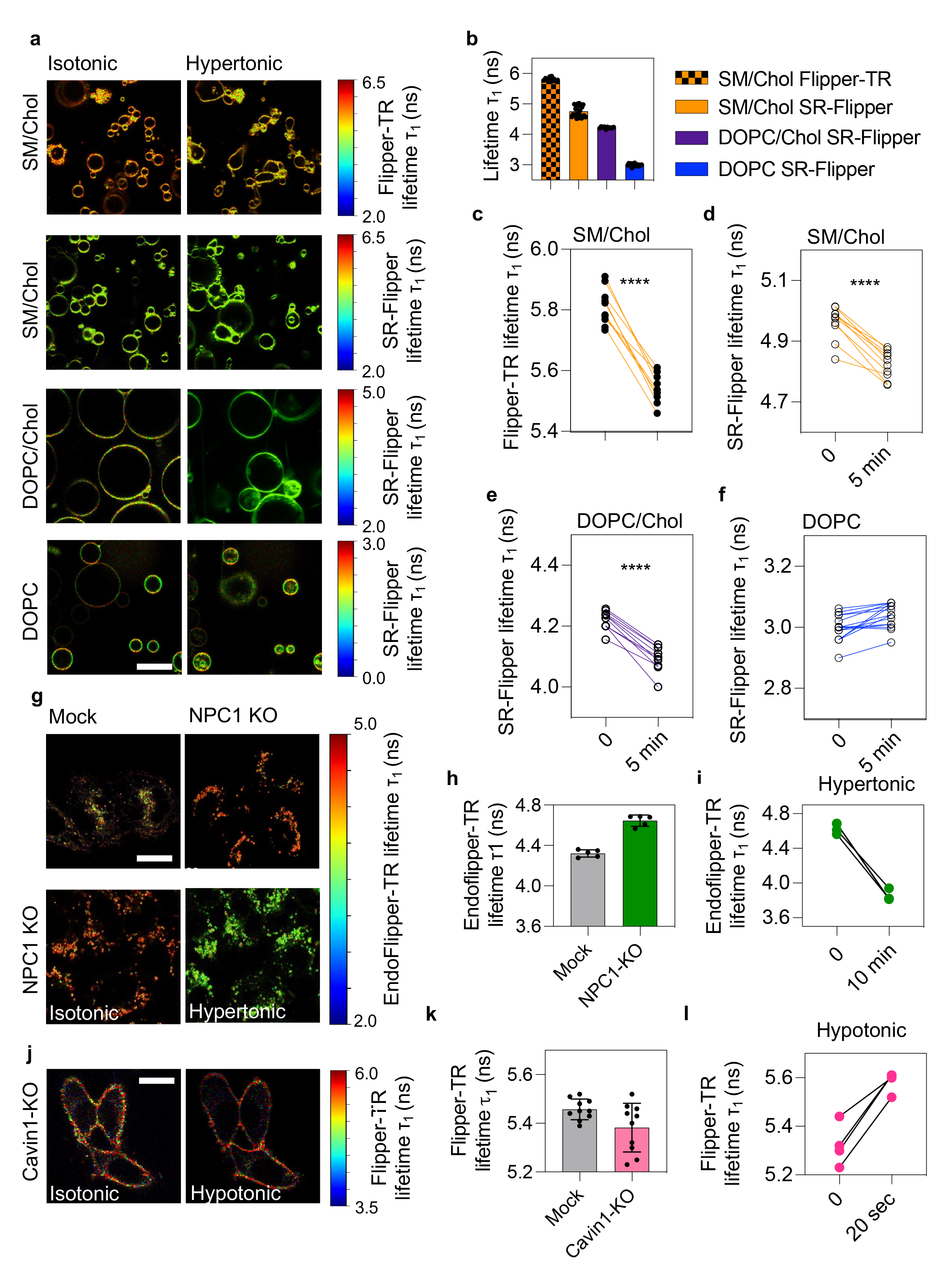
Flipper lifetime allow to detect membrane tension variations although if lipid composition is different. (a) Representative FLIM images before and after hypertonic shocks of SM/Chol containing GUVs stained with Flipper-TR or SR-Flipper, DOPC/Chol containing GUVs stained with SR-Flipper and of DOPC containing GUVs stained with SR-Flipper (d). (b) Graph showing the quantification of Flipper-TR and SR-Flipper lifetime (τ_1_, ns) before hypertonic shock on GUV of specified lipid composition. (c) Graph showing the quantification of Flipper-TR lifetime (τ_1_, ns) before and after hypertonic shock on SM/Chol GUV. (d-f) Graph showing the quantification of SR-Flipper lifetime (τ_1_, ns) before and after hypertonic shock on GUV of specified lipid composition. (g) Top : representative Endo Flipper FLIM images of HeLa MZ cells (left) and HeLa MZ NPC1 KO cells (right), which present accumulation of Cholesterol in endosomes. Bottom : representative Endo Flipper FLIM images of HeLa MZ NPC1 KO cells before hypertonic shock (left) and after hypertonic shock (right). (h) Graph showing the quantification of Endo Flipper lifetime (ns) of HeLa MZ cells and HeLa MZ NPC1 KO cells (i) Graph showing the quantification of Endo Flipper lifetime (τ_1_, ns) before and after hypertonic shock (j) Representative Flipper-TR FLIM images of HeLa Cavin1-KO cells before hypotonic shock (left) and after hypotonic shock (right). (j) Graph showing the quantification of Flipper-TR lifetime (τ_1_, ns) in HeLa cells (n = 10) and HeLa Cavin1-KO cells (n = 10) in isotonic medium. (k) Graph showing the quantification of Flipper-TR lifetime (τ_1_, ns) in HeLa Cavin1-KO cells (n = 4) after a hypotonic shock

Niemann-Pick C1 protein (NPC1) is responsible of the intracellular transport of Cholesterol and sphingolipids. Consequently, the depletion of NPC1 is responsible for cholesterol accumulation in endosomes. Consistently, LysoFlipper in HeLa MZ NPC1 KO cells have a higher lifetime than in WT HeLa cells indicating higher lipid packing (Fig 7g-h). Nonetheless, the probe is still able to report a change in membrane tension, as demonstrated by lifetime decrease under hypertonic treatment (Fig 7g,i).

The protein Cavin1 is a major component of caveolae and its deletion prevents caveolae assembly[10,44]. Interestingly, the absence of caveolae is associated with a smaller Flipper-TR lifetime, indicating either a lower membrane tension or a more disordered membrane composition (Fig 7j-k). Cavin1-KO cells show a lifetime increase upon hypotonic shocks (Fig 7j,i). Despite a potential difference of lipid composition, Flipper-TR is anyway able to report an increase of membrane tension.

## Conclusion

Here, we have reported the current technical difficulties we faced using Flipper probes, and how we overcame them. In the future, further developments of Flipper probes will allow us to bypass some of these problems. Technical improvements in the FLIM image acquisition and analysis may also help spread the use of Flipper probes to all biological samples of interest. Flipper probes remain an interesting tool for many applications, and in some cases, the only available tool to measure membrane tension.

## Acknowledgment

We thank Shankar Srinivas and Christophe Royer (University of Oxford) for isolating and staining the mouse embryos. FS is grateful for the support from the EMBO (ALTF 849-2020) and HFSP (LT000404/2021-L) long-term postdoctoral fellowships. FS thanks Scott Fraser (University of Southern California) for the support and supervision and acknowledges the Translational Imaging Center (University of Southern California) for access to instrumentation and expertise. A.C. also acknowledges funding from MCIU, PID2019-111096GA-I00; MCIU/AEI/FEDER MINECOG19/P66, RYC2018-024686-I, and Basque Government T1270-19. VD thanks Pierre-François Lenne (IBDM Marseille) for supervision and resources, and Katia Barrett for preparation of Xenopus explants. V.D. acknowledges support by an HFSP long-term postdoctoral fellowship (HFSP LT0058/2022-L) and the France-BioImaging infrastructure supported by the French National Research Agency (ANR–10–INBS- 04-01, Investments for the future). VD thanks Frank Schnorrer (IBDM Marseille) for access to the FLIM system. SM thank the University of Geneva, the National Centre of Competence in Research (NCCR) Chemical Biology (51NF40-185898), the NCCR Molecular Systems Engineering (51NF40-182895), and the Swiss NSF (Excellence Grant 200020 204175; SNSF-ERC Advanced Grant TIMEUP, TMAG-2_209190) for financial support. AR acknowledges funding from the Swiss National Fund for Research Grants N°310030_200793 and N°CRSII5_189996, the European Research Council Synergy Grant N° 951324 R2-TENSION.

## Materials and Methods

### Cell culture

HeLa-MZ cells (Lucas Pelkmans, University of Zurich), HeLa MZ-NPC1 KO cells, HeLa-Kyoto-smURFP, HeLa-Kyoto cells from Cell Lines Service (CLS) (330919) and Madin-Darby Canine Kidney II (MDCK-II) (ECACC, Cat. No. 00062107) were cultured in Dulbecco’s modified Eagle medium (DMEM 61965026, Thermo Fisher Scientific, Waltham, USA) supplemented with 10% fetal bovine serum (FBS 102701036, Thermo Fisher Scientific), 1% penicillin-streptomycin (15140122, Thermo Fisher Scientific) and 1% non-essential amino acids solution (NEAA 11140035, Thermo Fisher Scientific), at 37°C and 5% CO_2_. Cells were regularly tested negative for contamination with mycoplasma (Eurofins GATC Biotech, Konstanz, Germany). The HeLa ATCC Cavin1-KO cell line and corresponding control were cultured in DMEM, 4.5 g/L glucose supplemented in pyruvate. MDA MB 431 cells were maintained as described von Schwedler, U. K. et al. The protein network of HIV budding. Cell. The keratinocytes were grown in Keratinocyte-SF growth medium (INVITROGEN, 17005042) supplemented with 50 mg/mL bovine pituitary extract (BPE; Gibco) and 5 ng/mL prequalified human recombinant epidermal growth factor 1-53 (EGF 1-53; Gibco). Hippocampal neurons were grown in neuro basal medium. A549 and A596 were grown in RPMI 1640 Glutamax Medium (INVITROGEN 61870010). All cell media were supplemented with 10% foetal bovine serum (10270-106; Thermofischer) and 1% penicillin-streptomycin in a 5% CO_2_ incubator. Our HeLa-MZ cells were authenticated by Microsynth (Balgach, Switzerland), which revealed 100% identity to the DNA profile of the cell line HeLa (ATCC: CCL-2) and 100% identity over all 15 autosomal short tandem repeats (STRs) to Microsynth’s reference DNA profile of HeLa. Our cell lines are not on the list of commonly misidentified cell lines maintained by the International Cell Line Authentication Committee. Cells were authenticated by Microsynth (Balgach, Switzerland) and are mycoplasma negative, as tested by GATC Biotech (Konstanz, Germany), and are not on the list of commonly misidentified cell lines maintained by the International Cell Line Authentication Committee. A.Thaliana embryon was deposited inside a 2ml Eppendorf that contained a solution of 500ul of water with 2uM Flipper-TR final concentration and incubated 45 min at RT. Embryos were immobilized with 0.5% of low melting point agar for imaging. Alginate capsules and alginate tubes techniques were described respectively in [1]and [2].

### Reagents and drugs treatment

We obtained EZ-Link Sulfo-NHS-SS-Biotin, from Thermo Fisher Scientific. We obtained DOPC, DSPE-PEG-Biotin, DOPE, DOPS and lissamine rhodamine B sulfonyl (18:1) Rhod PE from Avanti Polar Lipids (Alabaster, AL). We obtained the different Flipper probes from Spirochrome (SC020) or from the lab of Prof. Stefan Matile. For the experiments involving blebbistatin, the concentrations of blebbistatin (SIGMA B0560-5MG) was equal to 10 uM and kept constant throughout the experiment. Cells were preincubated in DMEM with blebbistatin at 37 °C for 45 min. DMEM was replaced by Leibovitz with blebbistatin and Flipper-TR for 15 min. Blebbistatin was diluted in DMSO. Other reagents and chemicals were obtained from Sigma-Aldrich (St. Louis, MO).

### Live-cell FLIM imaging

For FLIM imaging, cells were seeded into 35mm MatTek glass bottom microwell dishes (MatTek Corporation). Cells were rinsed with 1ml Leibovitz’s medium (Thermofischer, 21083027) supplemented with 10% Foetal Bovine Serum and 1% Penicillin Streptomycin or Fluorobrite DMEM imaging medium (Life Technologies, Thermofisher) and incubated 15 min at 37°C with 1 μM Flipper (same concentration and incubation time for all Flippers). FLIM imaging was performed using a Nikon Eclipse Ti A1R microscope equipped with a Time Correlated Single-Photon Counting (TCSPC) module from PicoQuant. Excitation was performed using a pulsed 485nm laser (PicoQuant, LDH-D-C-485) operating at 20 MHz, and emission signal was collected through a bandpass 600/50nm filter using a gated PMA hybrid 40 detector or a TimeHarp 260 PICO implemented board (PicoQuant). For analysis, the SymPhoTime 64 software (PicoQuant) was used to fit the fluorescence decay data (from full images or selected pixels depending on the experiment) to a dual exponential reconvolution model, whereby the longer lifetime τ_1_ was extracted. Data are expressed as the mean ± s.d.

### Giant unilamellar vesicles (GUVs) electroformation

GUVs were produced as described[3] in water containing 250mM sucrose. Briefly, 20 μl of 1 mg/mL lipid mix (DOPC: DOPS, 6:4, mol:mol) were deposited on two indium tin oxide (ITO)-coated glass slides (70-100Ω resistivity, Sigma-Aldrich) and placed in a vacuum drying oven for 60’ for complete solvent evaporation. An O-ring of ∼1mm thickness was used as a non-leaky spacer between the two ITO slides, and the chamber was formed by compressing the two slides with spring metal tweezers. The formed chamber was filled with 400μl of 250mM sucrose solution (osmolarity adjusted to outside buffer solution, around 250 mOsm) and exposed to 1V AC-current (10Hz sinusoidal) at room temperature for 1 h. The resulting suspension was collected in a tube and used within the next days for experiments. Here, the lipid composition used were DOPC (99, 997%) and 0.003% DSPE-PEG(2000) Biotin, DOPC:Cholesterol at 70:30 Mol%, containing 0.003% DSPE-PEG(2000) Biotin or Brain Sphingomyelin:Cholesterol at 70:30 Mol% containing 0.003% DSPE-PEG(2000) Biotin.

### Hypertonic shock on GUVs labelled with Flipper or SR-Flipper

One channel of multi-well flow chambers (Ibidi) with 24×60mm glass coverslips (Menzel-Glaser) was coated with 0.1 mg/ml avidin in water for 10min, and rinsed 3X with water and 2X PBS. 2µl GUVs were added in 50µl PBS during 5min to facilitate avidin-biotin interactions, and then the solution was replaced by 40µl of 0.1mg/ml albumin-biotin in buffer 1 in order to saturate free avidin sites. The chamber was washed with 60µl of PBS and 1µM of specified Flipper was added on GUVs. For analysis, ROIs were selected by focusing on the GUV equatorial plane. For osmotic shocks, PBS was replaced with a pump at a rate = 60µl/min to avoid GUV damages by PBS supplemented with 250mM sucrose (final osmolarity of 500 mosm) or PBS diluted 2 times with water (125 mosm). A FLIM image was recorded before osmotic shock and 5min after the shock. Lifetime extraction was performed as previously described.

### Hypotonic shock on HeLa ATCC Cavin1-KO labelled with Flipper-TR

Cells were seeded into 35-mm MatTek glass-bottom microwell dishes. For hypotonic shocks, we diluted the cell imaging media with MilliQ water. Under the microscope, 1 mL of MilliQ water solution was added to the imaging dish containing 0.330 mL of isotonic buffer during imaging to reach a final osmolarity of 120 mOsm. Osmotic shocks were applied 10 seconds before the second timepoint.

### Micropipette aspiration on cells

The experimental set-up used to aspirate cells with a micropipette was the same as previously reported[3], and combines bright-field imaging, confocal scanning microscopy and FLIM imaging on an inverted Nikon Eclipse Ti microscope. Approximatively 1000 HeLa Cells in suspension were injected in the chamber (previously passivated with casein) containing complete medium supplemented with 0.5M of sucrose (in order to have a reasonably low plasma membrane tension which is necessary for pipette aspiration). A cell was aspirated by setting a negative pressure on the micropipette and FLIM images were acquired at different pressures. The analysis was performed as mentioned before.

### Fluorescence recovery after photobleaching (FRAP) experiments

Flipper was imaged using a ×100 1.4 numerical aperture oil DIC plan-apochromat VC objective (Nikon) with a Nikon A1 scanning confocal microscope. Photobleaching was performed in circular regions with four iterations of 488 nm at 100% laser intensity. For the analysis, the fluorescence background was subtracted from the intensity values of the ROIs. Those values were subsequently normalized to an unbleached area and the value of the intensity of the first frame after bleaching was subtracted. Finally, FRAP curves were normalized to the value of the pre-bleach fluorescence intensity. To determine the FRAP recovery kinetics, we fitted with the following double exponential function f(t) = α(1 − et − t0/τ1) + (1 − α)(1 − et − t0/τ2), where τ_1_ and τ_2_ are the time constants and α is the weight fraction of the exponential with characteristic time τ_1_. Diffusion coefficients were extracted with the formula: D=Rn^2^+Re^2^/8τ1/2 from[4].

### Experiments with GPMVs

Giant plasma membrane vesicles (GPMVs) were produced from HEK293T cells which were maintained under standard culture conditions, 37 °C, 5% CO2 with DMEM (Corning) 10 % FBS, Penicillin (100 U/mL) Streptomycin (100 µg/mL) (Lonza) and 2 mM L-Glutamine (Sigma Aldrich) following the procedure described previously[5,6]. Cells were seeded a day prior to the GPMV production and allowed to grow to a confluence of about 75 %. To induce vesiculation, growth media was removed, the cells washed once with GPMV buffer (containing 150 mM NaCl, 10 mM HEPES, 2 mM CaCl2 at pH 7.4), and next PFA and DTT were added to a final concentration of 25 mM and 5 mM, respectively. The cells were incubated for 2.5 h at 37 °C. The supernatant containing the GPMVs was harvested, and detached cells were removed by a short centrifugation (90 seconds at 1000 g). GPMVs were labelled with 1 µM Flipper-TR (final concentration) for 5 minutes before transfer to an imaging chamber (IBIDI 18well µ slide #1.5 glass). To non-specifically immobilize the GPMVs partially for imaging, the cover glass was treated with poly-D-Lysine (0.1 mg/mL) for 30 minutes and washed extensively with PBS as described previously[7].

### Lifetime imaging using Leica SP8 FALCON

One and two photon FLIM measurements were performed at a Leica SP8 equipped with FALCON and Dive modules (Leica Microsystems) and a 40x HC PL IRAPO 1.10 NA water-immersion objective (Leica Microsystems). Flipper fluorescence was excited using the white light laser (Leica Microsystems) with repetition rates of 80 MHz, 40 MHz, or 20 MHz and emission set to 488 nm in one photon mode or fluorescence was excited at 900 nm using a tunable Spectra Physics InSight X3 Ti:Saphire laser (pulsing at 80 MHz) in two photon mode. Fluorescence was collected between 500 nm and 700 nm on internal HyD-SMD detectors (in a descanned path for one photon and non-descanned path for two photon mode, respectively). The pinhole was set to 1.2 AU (at 550 nm). Images were acquired at 700 Hz scanning frequency at a zoom of 5 (512 by 512 pixels resulting in a pixel size of 113.9 nm/px). For one FLIM image 25 frames were accumulated to acquire at least 10^4^ photons at peak of the TCPSPC curve. Laser powers were adjusted when data were recorded at different repetitions rates to maintain similar count rates.

### Leica SP8 image processing

Image analysis and processing were performed in LAS-X (Leica Microsystems) and FIJI[8]. All lifetime data were obtained by applying a tailfit either for the whole image or on per pixel basis. For fitting Flipper decay data, a two-exponential model had to be employed. The shorter component was fixed based on the average short lifetime (of the whole image) for one respective dataset/condition (typically around 0.7 – 1.1 ns). The second component was extracted after fitting again (with short lifetime fixed) and presented as the tension sensitive component. In one photon mode, tail fitting was performed from 2.5 ns to 8.2 ns (80 MHz), 2.5 ns to 20 ns (40 MHz), or 2.5 ns 44 ns (20MHz). In two photon mode, fitting was performed from 2 ns to 12 ns (80 MHz). Generally, lower fitting boundary was about 0.5 ns from TCPSC maximum and on the edge of the estimated IRF. For the analysis of GUV and GPMV data, images were cropped to 320 by 320 pixels (36.4 µm by 36.4 µm), binned by a factor of 2 to increase photon count per pixel, and thresholded to exclude background pixels (for the GPMVs). Finally, the image fit was performed keeping the short lifetime component fixed and the longer component floating. Fitted lifetimes per image were exported to Excel. FLIM images were exported as raw images to .tiff (using 0.001 lifetime values per gray level accuracy). For presentation, images were thresholded using the intensity image and a rainbow LUT applied to the second lifetime component (all performed in FIJI as described previously[9]).

### Fluorescence Lifetime Imaging Microscopy of gastruloids and Xenopus explants

FLIM measurements were performed on a Zeiss LSM880 system (Carl Zeiss, Oberkochen, Germany) equipped with a time-resolved LSM upgrade (PicoQuant GmbH, Berlin, Germany) using a Plan-Apochromat 40x, 1.2NA water immersion objective. Images of 512×512 pixels per frame were acquired after excitation with a 510 nm pulsed laser diode operating at 20 MHz repetition rate. Fluorescence was split using a 560 nm dichroic mirror and detected by two SPAD detectors, after passing 550/49 nm and 600/50 nm bandpass filters, respectively. The pinhole was set to 1.4 AU. In each measurement, a minimum of 10^5^ photons at peak were recorded by accumulation of ca. 30 frames over a time period of 120 s. FLIM data were analysed using SymPhoTime64 software (PicoQuant GmbH, Berlin, Germany). A dual-exponential function was fitted to the decay curves recorded in each pixel, pooled from both detection channels. The short lifetime component was fixed to the average value (ca. 0.9 ns in Xenopus explants, ca. 1.5 ns in gastruloids) obtained by fitting a dual exponential function to the decay curve recorded on the whole image. The second, long-lifetime component was extracted to generate the presented FLIM Flipper tension maps.

### Two-photon microscopy

Two-photon imaging of gastruloids was performed on a Zeiss 510 NLO (Inverse - LSM) system (Carl Zeiss, Oberkochen, Germany) using a Plan-Apochromat 40x, 1.2NA water immersion objective. Fluorescence was excited using a femtosecond laser (Laser Mai Tai, DeepSee HP) and detected using two non-descanned GaAsP detectors (channel 1: 500-550 nm, channel 2: 575-640 nm). Per gastruloid, the same 2D plane was imaged at 800 nm to 1040 nm wavelength, in 20 nm intervals. To minimize sample movement, the imaging was performed at room temperature. The detected intensity values were normalized by the effective laser powers recorded using a power meter to correct for wavelength dependent variations of the laser output power.

### MDCK monolayers in an alginate tube

Alginate tubes were produced using a microfluidic device[2]. Briefly, 2.5% w/v sodium alginate (Protanal LF200S, FMC corporation, Philadelphia, USA), sorbitol (56755, Merck, Darmstadt, Germany) and a Matrigel (354234, Lot #5173011, Corning)/cell solution were injected in a 3D printed chip with three co-axial channels and the cell solution was incapsulated in the tubes by immersing the tip of the device in a gelation bath with 0.1 M calcium chloride (449709, Sigma-Aldrich, St Louis, USA), and pre-warmed to 37°C.

### MDCK monolayers in a PDMS roll

PDMS rolls were produced with a self-rolling thin bilayer[10]. Briefly, a toluene (T/2300/15, Fisher Chemical, New Jersey, USA)/PDMS (10 wt% of curing agent, Sylgard 184, Dow Corning, Michigan, USA) solution and a toluene/silicon oil (47 V 350, VWR, Pennsylvania, USA)/PDMS solution were successively spincoated, after reticulation (110°C, 30 min), on a fish gelatin-coated (G7041, Sigma Aldrich, Missouri, USA) PDMS substrate. After the silicon oil extraction in isopropanol (P/7500/15, Fisher Chemical) overnight, fibronectin (33016015, Gibco, New York, USA) was incubated at the surface to allow a cell adhesive surface. Once the monolayer was formed and the Flipper-TR probe was incubated, the PDMS bilayer was cut with a scalpel in the central region, producing two symmetric tubes and forcing the cell monolayer to roll.

### Gastruloid preparation

A complete description of the culture conditions and the protocol for making gastruloids is presented in Baillie-Johnson et al., 2015[11]. In the experiments presented here, gastruloids were generated from E14Tg2a.4 mouse embryonic stem cells (mESC) (MMRRC, University of California Davis, US) using an initial cell number of 100, and cultured according to the modified protocol described in Hashmi et al., 2022[12]. After 2 days of culture, gastruloids were incubated for 1h in culture medium supplemented with 1 µM Flipper-TR, and transferred to 35 mm culture dishes (MatTek Corp., Ashland, MA) for FLIM performed at 37°C, or two-photon imaging. Successful, homogeneous Flipper-TR staining of gastruloids was achieved at different time points of culture (tested and verified between 24h and 96h).

### Preparation of *Xenopus laevis* animal pole explants

Ovulation was stimulated in *Xenopus laevis* adult females by injection of Human Chorionic Gonadotropin (ChorulonR, 800 units/animal). On the following day, eggs were recovered by squeezing, fertilized *in vitro* with sperm from Nasco males, de-jellied in 2% cysteine hydrochloride (pH 8.0) and washed, first in water, then in 0.1X MBS (Modified Barth’s Saline). Embryos were kept in 0.1X MBS at 13°C or 18°C until they reached the desired stage. Animal cap explants were dissected at stage 8.5-9 as previously described[13] and kept in 0.5X MBS solution supplemented with 10 ng/ml Activin (RnD systems). Explants were transferred to a lipidure coated 35 mm culture dish (MatTek Corp., Ashland, MA), stained for 1h in medium supplemented with 1 µM Flipper-TR and stabilized for imaging using a viscous medium, 0.5X MBS supplemented with 1% methylcellulose. Due to the limited penetration of light into the specimen, FLIM was performed in the first cell layer. The experiments were performed at room temperature.

### Preparation of mouse embryo

Wildtype BALB/c mouse embryos at E6.5 were stained with 1 µM Flipper-TR (Spirochrome) for 2h as described in detail previously[9]. Imaging was performed at 37°C on a Leica SP8 equipped with a FALCON module. Fluorescence was excited using a 20x multi-immersion objective (Leica Microsystems) at 488 nm (white light laser, pulsing at 20 MHz) and recorded on the internal HyD-SMD detector (500-700 nm). Image size was 387.63 x 387.63 µm2. Images were processed in LAS-X (Leica Microsystems), thresholded to reject background photons, and binned by a factor of 6 to increase signal-to-noise ratio. The FLIM images were generated using the Phasor-FLIM analysis workflow. The Phasor data were median filtered (window of 5) and a rainbow look up table applied between 3.75 ns and 4.75 ns lifetime (phasor space). The image was exported as RGB for visualization purposes. Quantitative analysis should be performed on exported lifetime images.

## Notes

### Competing Interest Statement

The authors have declared no competing interest.

### Summary of Updates

Bibliography updated Small corrections done on Fig 3 Clarifications in the text

